# Maize *defective kernel5* is a bacterial tamB homolog required for chloroplast envelope biogenesis

**DOI:** 10.1101/372045

**Authors:** Junya Zhang, Shan Wu, Susan K. Boehlein, Donald R. McCarty, Gaoyuan Song, Justin W. Walley, Alan Myers, A. Mark Settles

**Affiliations:** Plant Molecular and Cellular Biology Program, University of Florida, Gainesville, FL; Horticultural Sciences Department, University of Florida, Gainesville, FL; Plant Pathology and Microbiology, Iowa State University, Ames, IA; Roy J. Carver Department of Biochemistry, Biophysics and Molecular Biology, Iowa State University, Ames, IA

**Keywords:** Chloroplast, Membrane biogenesis, Maize, Kernel, Membrane transporter, Metabolism

## Abstract

Chloroplasts are of prokaryotic origin with a double membrane envelope that separates plastid metabolism from the cytosol. Envelope membrane proteins integrate the chloroplast with the cell, but the biogenesis of the envelope membrane remains elusive. We show that the maize *defective kernel5* (*dek5*) locus is critical for plastid membrane biogenesis. Amyloplasts and chloroplasts are larger and reduced in number in *dek5* with multiple ultrastructural defects. We show that *dek5* encodes a protein homologous to rice *SUBSTANDARD STARCH GRAIN4* (*SSG4*) and *E.coli* tamB. TamB functions in bacterial outer membrane biogenesis. The DEK5 protein is localized to the chloroplast envelope with a topology analogous to TamB. Increased levels of soluble sugars in *dek5* developing endosperm and elevated osmotic pressure in mutant leaf cells suggest defective intracellular solute transport. Both proteomics and antibody-based analyses show that *dek5* chloroplasts have reduced levels of chloroplast envelope transporters. Moreover, *dek5* chloroplasts reduce inorganic phosphate uptake with at least an 80% reduction relative to normal chloroplasts. These data suggest that DEK5 functions in plastid envelope biogenesis to enable metabolite transport.

## INTRODUCTION

Plastids are essential organelles for plants. Higher plants differentiate specialized plastids distinguished by structure, pigmentation, and function, such as photosynthetically active chloroplasts in leaf tissue and starch accumulating amyloplasts in the cereal endosperm (Jarvis and López-Juez, 2013). Plastids originated through endosymbiosis approximately 1.5 billion years ago when cyanobacteria were acquired into eukaryotic host cells (Yoon et al., 2004). Extant cyanobacteria are Gram-negative with inner and outer plasma membranes. Chloroplasts have a similar double membrane structure with the inner and outer envelope membranes likely corresponding to the membranes of the bacterial endosymbiont (Gould et al., 2008; Gross and Bhattacharya, 2009). The vast majority of chloroplast proteins are nuclear-encoded and synthesized as precursor proteins on cytosolic ribosomes for post-translational import into plastids (Jarvis, 2008). Nuclear-encoded precursors are imported through the TOC/TIC translocons located on the chloroplast envelope membranes (Keegstra and Cline, 1999; Cline and Dabney-Smith, 2008).

Plant cellular metabolism pathways integrate enzymes located in different subcellular organelles with the plastid having a major role in primary metabolism (Bowsher and Tobin, 2001). Transport of solutes and metabolites across chloroplast envelope membranes is important to integrate chloroplast metabolism with the cytosol and other cellular organelles. Chloroplast envelopes exchange ions, carbohydrates, nucleotides, and amino acids to support metabolic pathways in which the chloroplast has unique enzymatic activities (Block et al., 2007; Facchinelli and Weber, 2011).

The inner envelope of the chloroplast has multiple solute translocators and is known to be the major permeability barrier for metabolites (Flügge, 1999; Fischer, 2011). Inner envelope translocators are integral membrane proteins with two pathways for insertion. During protein import, some inner envelope membrane proteins are transferred to the membrane through a stop-transfer mechanism. Other inner envelope membrane proteins complete import into the stroma and are inserted into the membrane similar to post-translational translocation of secreted bacterial proteins (Lubeck et al., 1997; Nada and Soll, 2004).

The outer envelope is thought to be permeable to solutes with a molecular weight of less than 10 kDa, which is similar to the outer membrane of Gram-negative bacteria (Flügge and Benz, 1984). Porins are integral membrane channels that facilitate this nonspecific diffusion of small solutes in Gram-negative bacteria (Nikaido, 1994). Many chloroplast outer envelope proteins (OEP) have a β-barrel structure similar to porins and were hypothesized to also facilitate nonspecific diffusion; however, biochemical analysis shows more selective transport. Pea OEP21 transports inorganic phosphate (Pi), triose phosphates, and 3-phosphoglycerates (Hemmler et al., 2006). OEP24 allows diffusion only of triose phosphates, dicarboxylic acids, charged amino acids, ATP, and Pi (Pohlmeyer et al., 1998). OEP40 is only permeable to glucose, glucose-1-phosphate, and glucose-6-phosphate (Harsman et al., 2016). OEP16 and OEP37 are selective for amino acids and peptides and even have tissue specific expression patterns (Pohlmeyer et al., 1997; Goetze et al., 2006; Pudelski et al., 2012). Thus, OEP channels studied so far show specificity for distinct metabolites challenging the notion that the outer envelope is a nonspecific molecular sieve.

Relatively little is known about the biogenesis pathways of β-barrel OEPs. In Gram-negative bacteria, most β-barrel outer membrane proteins require the β-barrel assembly machinery (BAM) for their correct folding and function (Hagan et al., 2011; Selkrig et al., 2014). The translocation and assembly module (TAM) is also important for bacterial outer membrane biogenesis and extracellular secretion. TAM is composed of the TamA and TamB proteins with TamA localized to the outer membrane and TamB localized to the inner membrane (Selkrig et al., 2012). Mutations of TAM proteins in different bacterial species can alter membrane morphology or block secretion of some toxins into media (Selkrig et al., 2012; Shen et al., 2014; Iqbal et al., 2016). Phylogenetic analysis showed that TamA is restricted to *Proteobacteria*; whereas, TamB is widely distributed across most Gram-negative bacterial lineages (Heinz et al., 2015). In *Borrelia burgdorferi*, a spirochete that does not encode TamA, the TamB ortholog interacts with a BAM subunit supporting the idea that TamB can participate in assembly of β-barrel proteins through multiple outer membrane protein assembly machineries (Iqbal et al., 2016).

Here, we show that the maize *defective kernel5* (*dek5*) mutant has plastid division defects that disrupt endosperm starch accumulation. Mutant *dek5* seedling leaves have fewer and larger chloroplasts with defects in chloroplast membranes. Molecular identification of the *dek5* locus demonstrated that it encodes a predicted TamB homolog. Contrary to a prior report for the rice DEK5 ortholog (Matsushima et al, 2014), the maize DEK5 protein is localized to the chloroplast envelope with analogous topology to TamB. The *dek5* mutant has altered envelope ultrastructure, reduces outer envelope protein accumulation, alters inner envelope protein levels, and blocks Pi uptake. These data suggest that *Dek5* plays an important role in plastid envelope biogenesis and illustrate the importance of selective solute transport across the plastid envelope.

## RESULTS

### Starch accumulation defects in *dek5* kernels

The Maize Genetics Cooperative Stock Center maintains six mutant alleles of the recessive *dek5* locus: *dek5-N874A*, *dek5-N961,* and *dek5-N1339A* isolated from EMS-mutagenesis; *dek5-PS25* and *dek5-MS33* isolated from a *Robertson’s Mutator* transposon-tagging population, and the spontaneous *dek5-Briggs* allele isolated at a commercial breeding company (Neuffer and Sheridan, 1980; Scanlon et al., 1994; Sachs, 2009). At maturity, each allele conditions shrunken or collapsed endosperm, while embryo development is frequently normal (Figure 1A-B, Supplemental Figure 1A). This visual phenotype resembles that of *brittle1* (*bt1*), *brittle2* (*bt2*), and *shrunken2* (*sh2*) mutants blocked early in starch biosynthesis (Supplemental Figure 1B). Among the alleles, *dek5-Briggs* and *dek5-N1339A* have more severe phenotypes (Supplemental Figure 1A). Unless noted, *dek5-MS33* was used for phenotype analysis.

**Figure 1.**
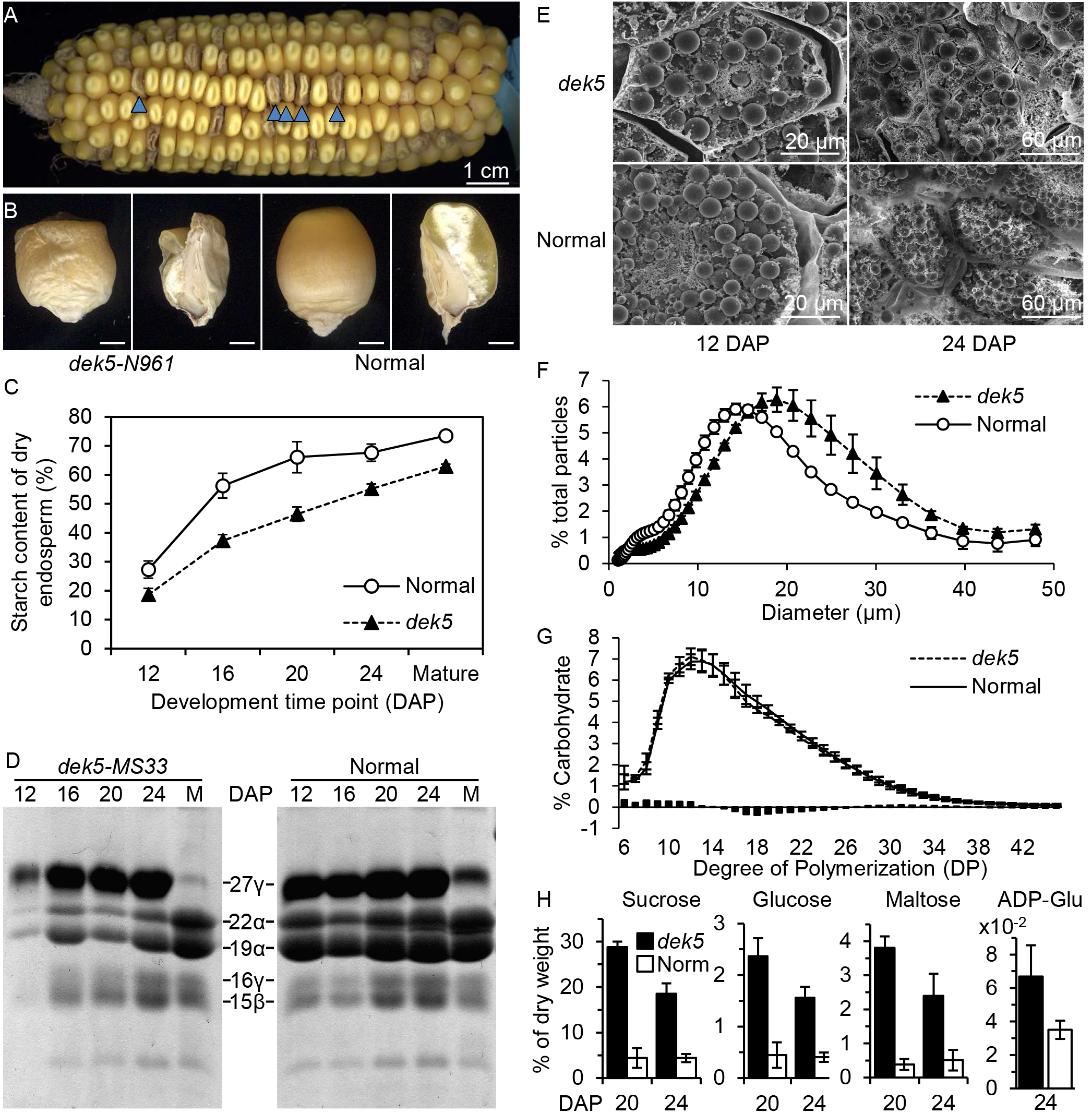
*dek5* kernel phenotypes. (A) Ear segregating for *dek5-N961* in the B73 genetic background. Blue arrowheads point to *dek5* kernels in a single row. (B) Abgerminal view and sagittal sections of *dek5-MS33* and normal sibling. Scale bars are 2 mm. (C) Endosperm starch for *dek5-MS33* and normal siblings. Mean and standard deviation of three biological replicates are plotted. (D) Zein levels during development and at maturity (M). Protein was loaded on equal dry tissue weight basis. (E) SEM of starch granules in *dek5-MS33* and normal sibling endosperm. (F) Distribution of mature endosperm starch granule size from *dek5-MS33* and normal siblings. Particle sizes <1 μm and >50 μm are excluded. The plot accounts for 86% of total particles. Mean and standard deviation of four biological replicates are plotted. (G) Amylopectin chain-length distribution from *dek5-MS33* and normal siblings. Mean and standard deviation are plotted for three biological replicates. Bars plot the difference of normal minus mutant. (D) Metabolite analysis of developing *dek5-MS33* (black) and normal (white) sibling endosperm tissue at 20 and 24 DAP. Mean and standard deviation are plotted for three biological replicates.

Single-kernel near infrared reflectance (NIR) spectroscopy analyses indicated *dek5* mutants have reduced starch and increased oil content compared to normal siblings (Supplemental Table 1). The *dek5* starch deficiency is apparent in developing endosperm at 12 days after pollination (DAP) and extending to maturity (Figure 1C). Relative zein content, particularly α-zeins, is also reduced in *dek5* mutants (Figure 1D). Both *bt1* and *bt2* have reduced zein content associated with reduced starch (Lee and Tsai, 1984). Non-zein protein content had no obvious changes when estimated by SDS-PAGE (Supplemental Figure 2A).

Hand sections of developing kernels had uneven iodine staining of starch in *dek5* endosperm from 16-24 DAP, even though the mutant kernels enlarged to a similar extent compared to normal siblings (Supplemental Figure 2B). Like *bt1*, *bt2*, and *sh2* starch biosynthesis mutants, the *dek5* collapsed endosperm phenotype only becomes apparent as mature kernels dry down. Endosperm starch granules have a normal diameter at 12 DAP but increase significantly in *dek5* at 24 DAP (Figure 1E, Supplemental Figure 2C). Mature endosperm granule size was measured by particle analysis with a 20% increased average size in *dek5* (Figure 1F). Reduced total starch concomitant with increased granule size implies *dek5* endosperm accumulates fewer granules than normal.

The average chain length distribution of the amylopectin component of endosperm starch was determined after enzymatically converting the branched polymer to a population of linear α(1→4)-linked glucan chains. The frequency of each length from 6 to 44 glucosyl units is essentially identical in *dek5* and normal siblings (Figure 1G). Despite the enlarged granule size, there is no apparent change in the net activities of starch synthases and starch branching enzymes to assemble semi-crystalline amylopectin. However, soluble sugar levels are consistent with reduced accumulation of starch. At 20 and 24 DAP, *dek5* endosperm has elevated levels of sucrose, glucose, and maltose as well as higher levels of ADP-glucose at 24 DAP (Figure 1H). Sucrose, glucose, and ADP-glucose are soluble precursors of starch. On a fresh weight basis, sucrose and glucose are elevated similarly in *dek5*, *sh2*, and *bt1* with *dek5* sucrose and glucose being 4.2-fold and 3.4-fold higher than normal siblings, respectively (Tobias et al., 1992). The higher levels of ADP-glucose in *dek5* are similar to those observed for *sh2; bt1* double mutants (Shannon et al., 1996).

### Chloroplast defects in *dek5* plants

Depending upon the allele, 5-40% of *dek5* kernels will germinate and develop pale green seedlings with non-clonal, white variegation. Mutant seedlings are stunted at 7 days after sowing (DAS) and usually die in 2-3 weeks (Figure 2A, Supplemental Figure 1A). The Mo17 and W22 genetic backgrounds enhance *dek5* resulting in a nearly empty pericarp (*ep*) kernel phenotype, lower germination, and more severe seedling phenotypes. The *dek5-Briggs* allele was the most severely modified (Supplemental Figure 3A). By contrast, introgression of *dek5* into B73 improved grain-fill, germination, and mitigated seedling lethality (Supplemental Figure 3B). The mild *dek5-MS33* allele could complete a life cycle to produce all mutant kernels (Supplemental Figure 3C).

**Figure 2.**
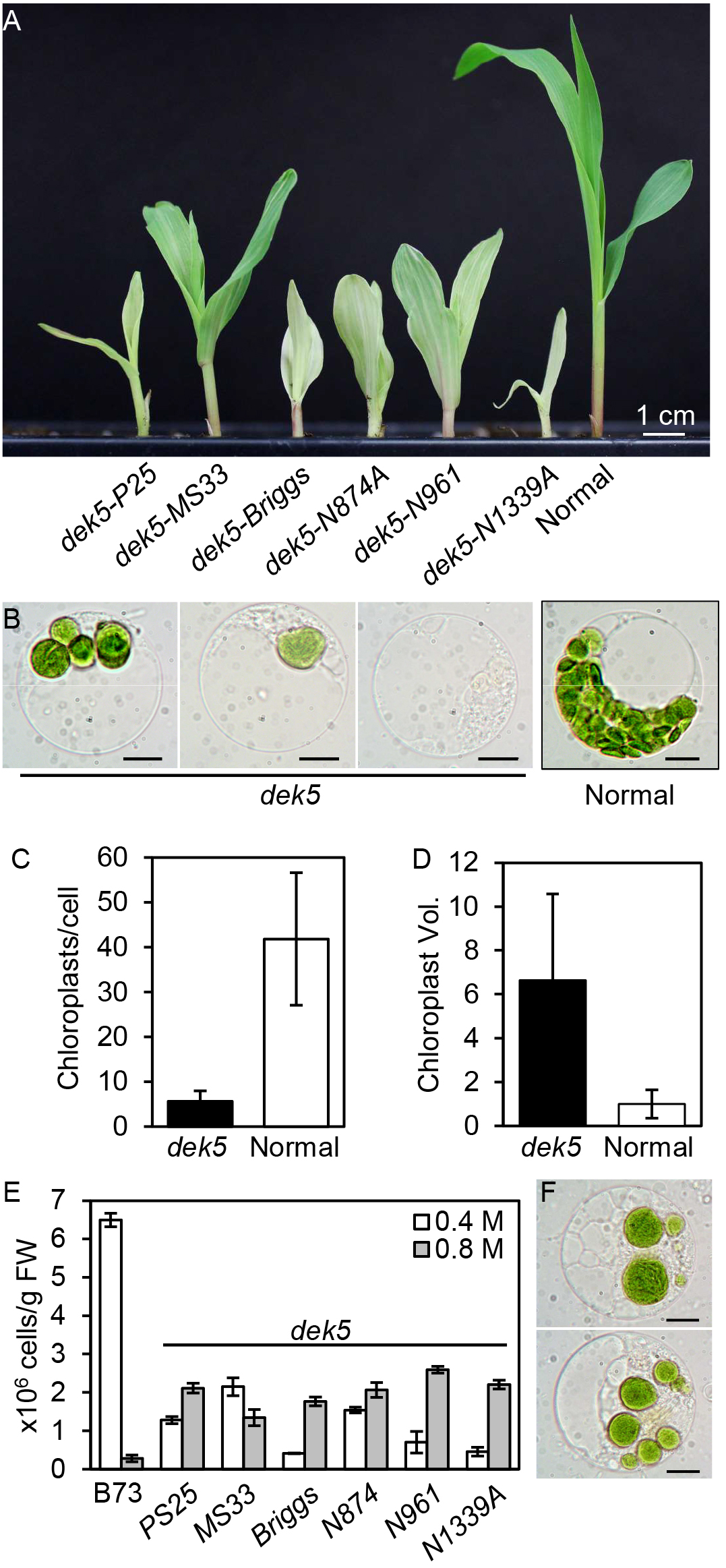
Fewer and larger chloroplasts in *dek5* seedling leaf cells. (A) Representative *dek5* seedlings for six alleles. Scale bar is 1 cm. (B) Example protoplasts from *dek5-MS33* and a normal sibling. Scale bar is 10 μm. (C-D) Summary statistics from LCSM Z-stack imaging of at least 60 *dek5-MS33* and normal sibling protoplasts showing chloroplasts per cell (C) and relative chloroplast volume (D). Error bars are standard deviation. (E) Protoplast yield with increasing mannitol. Mean and standard errors are plotted for three technical replicates. (G) *dek5-MS33* mutant protoplasts contain multiple vacuole compartments. Scale bar is 10 μm.

The pale green, variegated seedling leaves in *dek5* are due to multiple chloroplast abnormalities. Photosynthetic pigments and subunits of the electron transport chain are reduced (Supplemental Table 2; Supplemental Figure 4A). Seedling leaf protoplasts showed *dek5* mutants have larger and fewer chloroplasts compared to normal siblings with a subset of mutant protoplasts having one chloroplast or no chloroplasts (Figure 3B). Z-stack imaging of chlorophyll florescence showed *dek5* protoplasts have 7.5-fold fewer chloroplasts with a 6.6-fold increase in chloroplast volume (Figure 2C-D, Supplemental movie 1).

**Figure 3.**
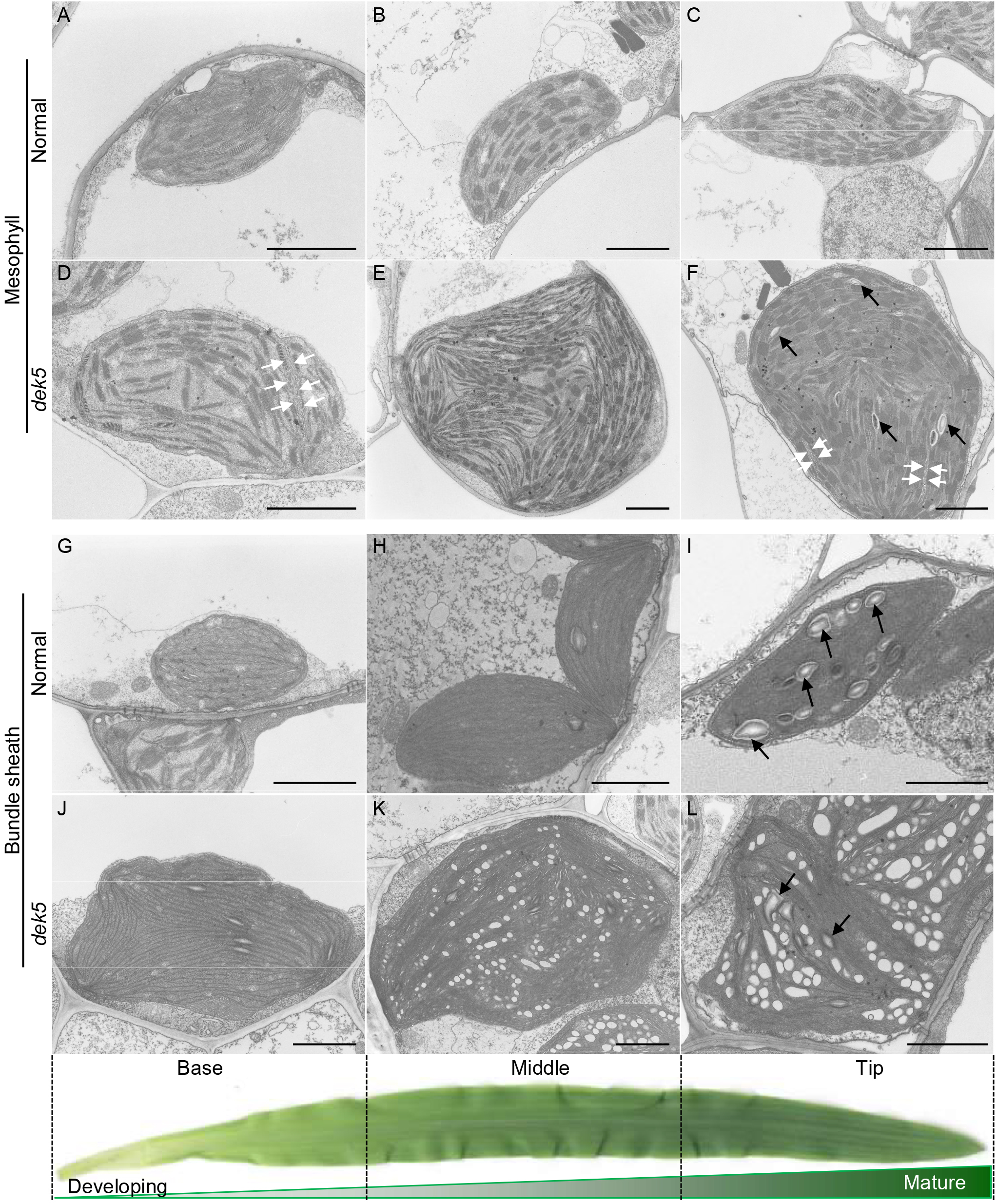
TEM analysis of dek5 and normal sibling leaf chloroplasts. (A-C) Normal mesophyll cell chloroplasts. (D-F) Mutant *dek5-MS33* mesophyll cell chloroplasts. White arrows indicate envelope ingrowths. Black arrows indicate starch granules. (G-I) Normal bundle sheath cell chloroplasts. (J-L) Mutant bundle sheath cell chloroplasts. Scale bar is 2 μm in all panels. Black arrows indicate starch granules.

Mutant protoplasts isolated in standard buffers with 0.4 M mannitol had a low recovery rate, while parallel isolations using 0.8 M mannitol increased recovery nearly 2-fold on average (Figure 3F, Student’s t-test p=0.02). The *dek5* cells isolated in high osmotic buffer frequently contained multiple vacuolar compartments, which is a morphological characteristic of autophagy or programed cell death (Figure 3G) (van Doorn et al., 2011). These data suggest that *dek5* cells are more likely to lyse with standard buffers and have an altered osmotic pressure.

Transmission electron microscopy (TEM) revealed multiple ultrastructural defects in *dek5* chloroplasts. Chloroplasts in expanding maize seedling leaves develop in a gradient. At the base, differentiating plastids have few internal membranes, while chloroplasts at the middle and tip of the leaf have developed thylakoids (Pogson et al., 2015). Maize is also a C4 plant with both mesophyll and bundle sheath cells developing distinct chloroplast morphology. Sections from the base, middle, and distal tip of *dek5* seedling leaves had mesophyll and bundle sheath chloroplasts with developed internal membranes, although chloroplasts were much larger (Figure 3). Consistent with protoplast Z-stack imaging, *dek5* mesophyll chloroplasts have 4-fold larger cross-sectional area than normal siblings (Figure 4A).

**Figure 4.**
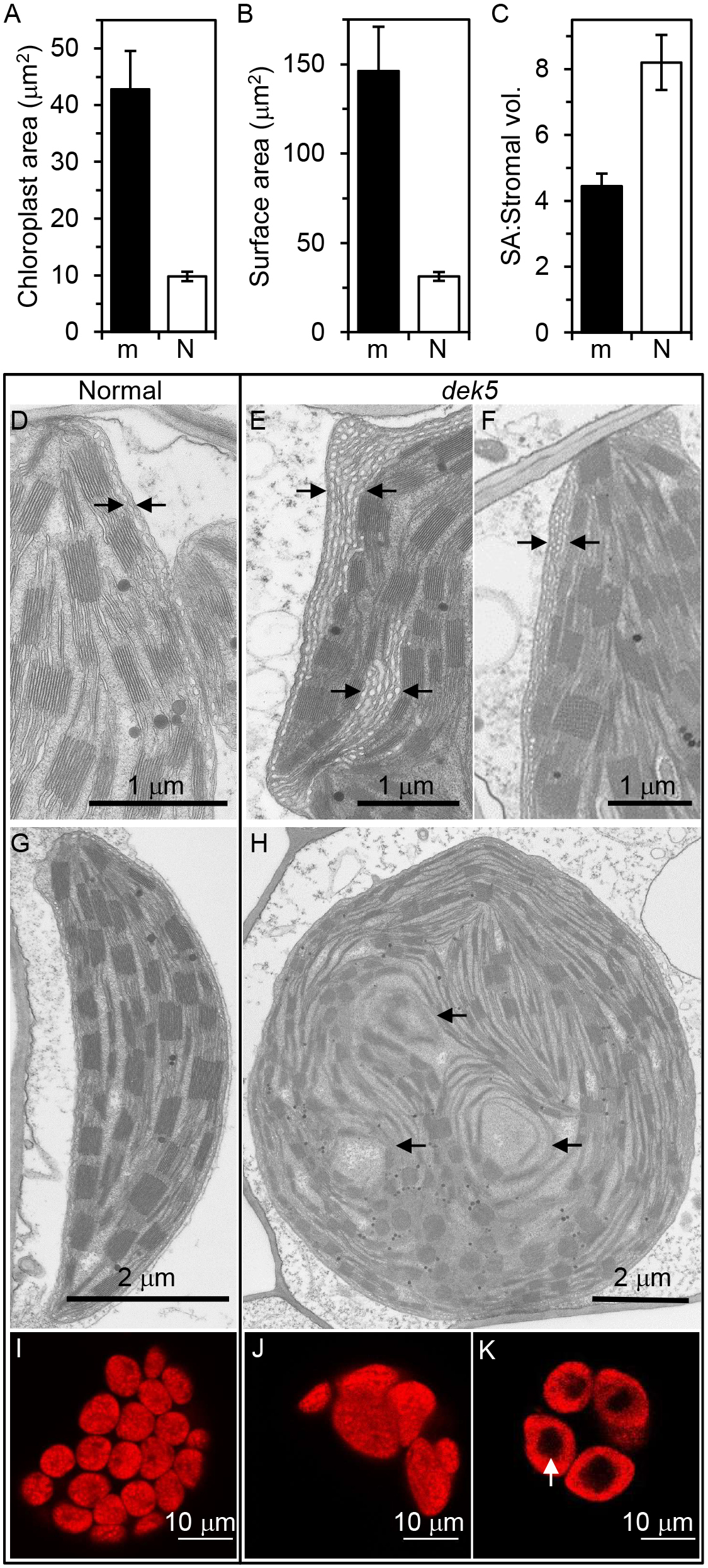
Range of *dek5* chloroplast membrane defects. (A-C) Quantitative chloroplast traits from TEM images. Each plot shows mean and standard error of at least 30 chloroplasts from seven *dek5* (m) and three normal (N) sibling leaves. (A) Chloroplast cross-section area measured by ImageJ. (B) Chloroplast surface area modeled as an ellipsoid from the long and short axis. (C) Surface area (SA) to stromal volume ratio. (D) Normal chloroplast envelope (arrows). (E-F) Thick, chloroplast reticulum (arrows) in *dek5-MS33*. (G) Normal mesophyll chloroplast. (H) Disorganized thylakoids in *dek5-MS33*. (I-K) Individual LSCM cross-sections showing protoplast chlorophyll autoflorescence. (I) Normal sibling with red signal throughout each chloroplast. (J) A *dek5-MS33* protoplast with fully fluorescent chloroplasts. (K) Reduced chlorophyll fluorescence in the central thylakoids.

Many *dek5* mesophyll chloroplasts show similar thylakoid development to normal siblings (Figure 3D-F). There are also multiple thylakoid centers and a more spherical organelle shape in *dek5* leaves (Figure 3E-F). Normal chloroplasts have two thylakoid centers defining the long axis of the organelle (Figure 3B-C).

C4 plants preferentially localize starch biosynthetic enzymes and starch granules in bundle sheath chloroplasts (Majeran et al., 2005; Leegood, 2008; Weise et al., 2011). Large starch granules were observed in normal bundle sheath chloroplasts at the tip of the leaf (Figure 3I). Like *dek5* mesophyll chloroplasts, mutant bundle sheath chloroplasts are much larger; however, these contained many vesicular structures with a few, small starch granules (Figure 3L). Some *dek5* mesophyll chloroplasts also had small starch granules, which were not observed in normal siblings (Figure 3F). These observations suggest *dek5* has defects in regulating starch biosynthesis or transporting fixed carbon to bundle sheath cells.

The spherical, larger *dek5* chloroplasts are expected to alter the ratio of envelope, stroma, and thylakoid. We found that the cross-sectional area of the chloroplast could be modeled as an ellipse from long and short axis measurements, suggesting that the entire chloroplast could be modeled as an ellipsoid (Supplemental figure 5A-B). These estimates indicate a 14.7-fold increase in chloroplast volume and an 8.6-fold increase in stromal volume of *dek5* mutants (Supplemental figure 5C). However, envelope surface area is only increased by 4.7-fold (Figure 4B). Thus, the surface area to volume ratio of *dek5* chloroplasts or stroma is reduced by 2-fold relative to normal chloroplasts indicating a relative reduction in envelope biogenesis (Figure 4C; Supplemental figure 5C).

TEM also revealed a range of *dek5* chloroplast membrane defects. Mutant chloroplasts had unusual envelope ingrowths as well as an expanded peripheral reticulum at the envelope (Figure 4D-F) (Szczepanik and Sowiński, 2014). Although thylakoid grana have a relatively normal appearance, they are 40% larger in diameter in *dek5,* and internal thylakoid membranes appeared disorganized in some chloroplasts (Figure 4G-H; Supplemental Figure 4B-F). In some cases, internal vesicles and completely disorganized plastids were observed (Supplemental Figure 4F-G). These phenotypes were observed both in mesophyll and bundle sheath cells immediately adjacent to cells with intact chloroplasts suggesting that the defective membranes are unlikely to be an artifact from fixation or sample processing. Mutant chloroplasts with a loss of internal chlorophyll autoflorescence were also observed by LSCM in 16 of 51 *dek5* protoplasts that had full Z-stack imaging (Figure 4I-K, Supplemental movie 1). These data indicate that *dek5* has pleiotropic plastid membrane biogenesis defects.

### *Dek5* is homologous to *tamB*

The *dek5* locus was mapped to the short arm of chromosome 3 using B-A translocations (Neuffer and Sheridan, 1980). We fine-mapped the *dek5-PS25* allele to a genomic interval of 460 kbp containing 13 gene models in the B73 RefGen_v3 genome assembly (Figure 5A). Mu-Seq analysis of *Mutator* transposon insertions in the *dek5-PS25* and *dek5-MS33* identified an insertion within the fine-map interval in GRMZM2G083374. Co-segregation of the *Mu* insertion site with the *dek5* seed phenotype was confirmed, and two additional *Mu* insertion alleles from the UniformMu reverse genetic resource failed to complement *dek5-PS25* (Supplemental Figure 6A-B). There is a *Mu8-*like transposon in *dek5-PS25* and a *Mu1* insertion in *dek5-MS33* at the identical position in exon 1 indicating these are independent alleles. The *dek5-Briggs* allele has a frameshift mutation with a single base insertion at exon 10 that results in a downstream pre-mature termination codon. Molecular identification of five non-complementing alleles indicates GRMZM2G083374 is the *dek5* locus (Figure 5B).

**Figure 5.**
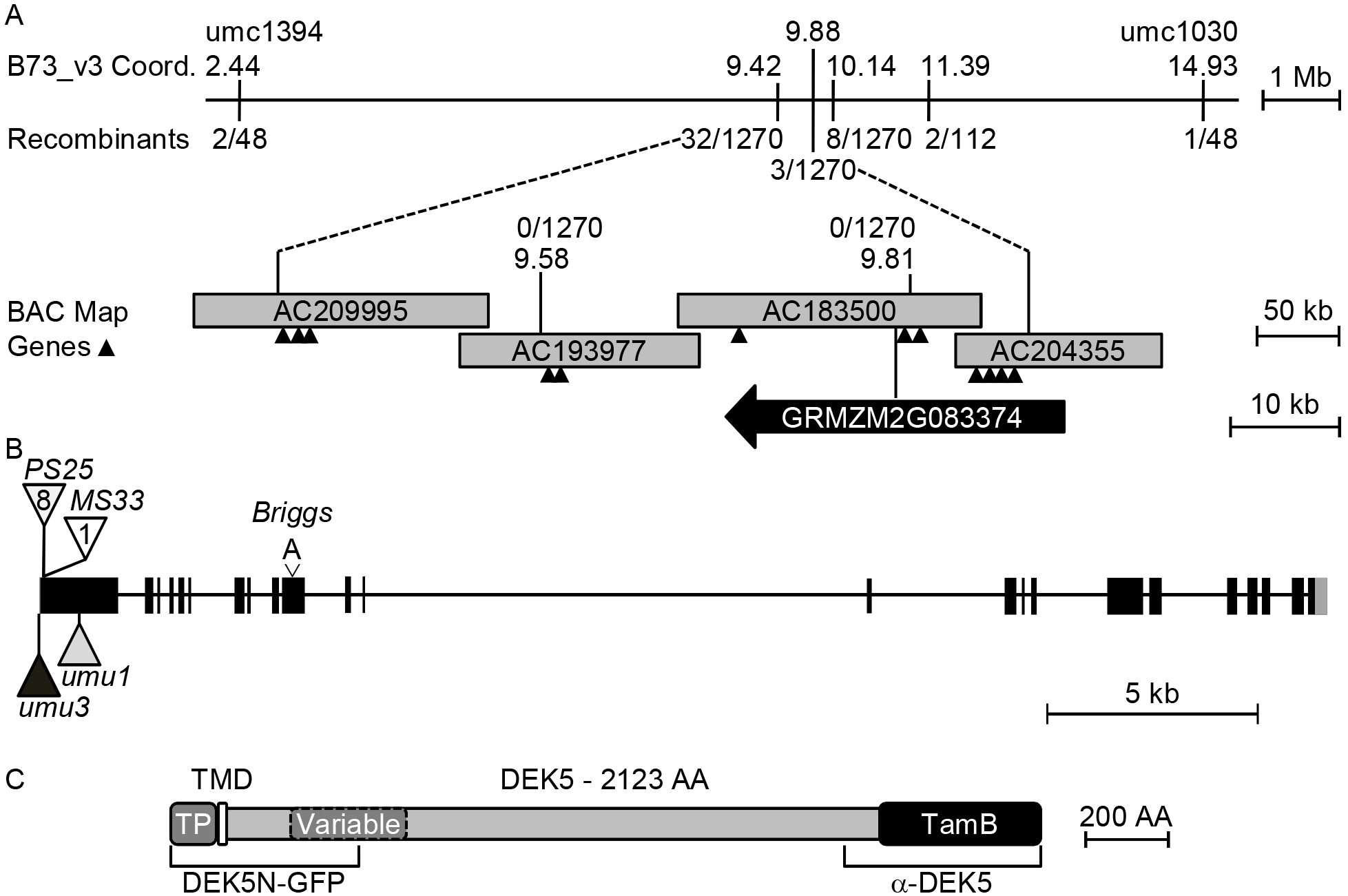
Map-based cloning of the *Dek5* gene. (A) Integrated physical-genetic map of the *dek5* locus. B73 AGP_v3 physical coordinates are given for molecular markers with recombinants/number of meiotic products genotyped. Bacterial artificial chromosome (BAC) sequences were aligned with blastn to determine overlap regions. Genes in the fine-map interval are denoted with triangles except for *dek5*, GRMZM2G083374. (B) *Dek5* locus schematic. Black boxes are coding exons, grey boxes are untranslated regions, and introns are lines. Triangles indicate *Mu* insertions with *dek5-PS25* having a *Mu8-like* insertion at the same site as *dek5-MS33* with a *Mu1* insertion. (C) DEK5 protein domains: N-terminal transit peptide (TP), a single-pass α-helical transmembrane domain (TMD), and a C-terminal TamB domain (DUF490). Brackets indicate protein regions used for a C-terminal GFP fusion (DEK5N-GFP) and polyclonal antibody production (α-DEK5).

The *dek5* genomic locus is predicted to be a single-copy gene that is >30 kbp. We amplified and sequenced the B73 cDNA to experimentally validate the predicted gene model. The sequenced *Dek5* mRNA has a different structure than GRMZM2G083374. It consists of 23 exons with a genomic transcriptional unit of 30,833 bp (Figure 5B). RT-PCR primers spanning exons 10-11 confirmed *Dek5* expression during kernel, seedling, and reproductive development (Supplemental Figure 6C).

*Dek5* encodes a predicted protein of 232.9 kDa that has an N-terminal chloroplast transit peptide (TP), a transmembrane domain (TMD), and a C-terminal TamB domain, formerly DUF490 (Figure 5C). In rice, a missense mutation in the *Dek5* ortholog, *SUBSTANDARD STARCH GRAIN4* (*SSG4*) results in enlarged starch granules and chloroplasts similar to *dek5* (Matsushima et al., 2014). Both the maize and rice domain structure is similar to the *E. coli* TamB protein (Selkrig et al., 2012). However, phylogenetic analysis indicates that DEK5 has additional conserved sequence domains within land plants (Matsushima et al., 2014; Supplemental Figure 7).

### DEK5 localizes to the chloroplast envelope

Chloroplast targeting of the DEK5 N-terminal TP and predicted TMD was tested by fusing the 5’ 1,374 bp of the *Dek5* open reading frame (ORF) with *EGFP* (Figure 5C, Supplemental Figure 8). Transient expression of DEK5N-GFP in *Nicotiana benthamiana* leaves resulted in a punctate ring of GFP signal surrounding red chlorophyll fluorescence from thylakoid membranes (Figure 6B). This localization pattern is identical to known chloroplast envelope membrane proteins, such as CHUP1, OEP7, Tic40, and Tic20 (Oikawa et al., 2008; Machettira et al., 2011; Breuers et al., 2012). However, it contrasts with the stromal subcellular localization reported for an N-terminal SSG4-GFP fusion protein from rice that only encoded part of the conserved N-terminal domain of DEK5 orthologs (Supplemental Figure 8) (Matsushima et al., 2014).

**Figure 6.**
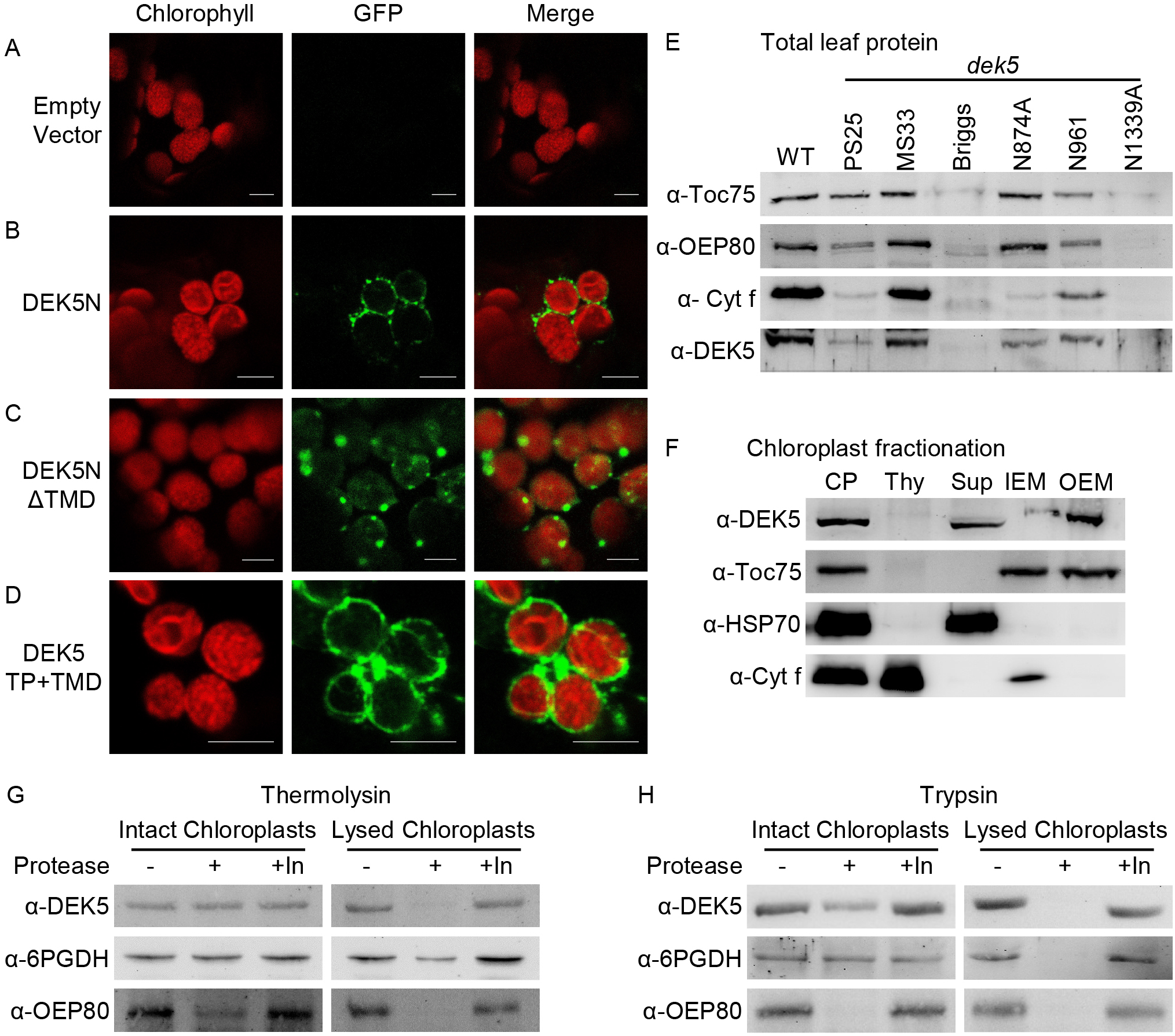
DEK5 topology. (A-D) Confocal micrographs after transient expression of GFP fusion proteins in *N. benthamiana* leaves: (A) Empty vector is pB7FWG2; (B) DEK5N-GFP has the N-terminal region shown in Fig. 5C; (C) DEK5NΔTMD-GFP has an in-frame deletion of the predicted transmembrane domain (TMD); (D) DEK5TP+TMD-GFP includes on the predicted chloroplast targeting sequence and TMD. Scale bar is 5 μm in all panels. (E) Immunoblots of total leaf protein with outer envelope Toc75 and OEP80, thylakoid membrane Cyt f, and DEK5 proteins. (F) Immunoblots of normal chloroplast subfractionation with whole chloroplasts (CP), thylakoid (Thy), soluble (Sup), inner envelope membrane (IEM), and outer envelope membrane (OEM) fractions. HSP70 is a soluble, stromal protein. (G-H) Protease protection assay with purified chloroplasts treated with either thermolysin (G) or trypsin (H) proteases (+) or the protease plus protease inhibitors (+In).

Structure-function analysis mapped the DEK5 envelope localization domain to the N-terminal TMD. Deletion of the predicted TMD sequence from DEK5N-GFP resulted in DEK5NΔTMD-GPF showing punctate distribution of GFP fluorescence overlapping with chlorophyll fluorescence indicating targeting within plastids (Figure 6C). Fusing only the DEK5 TP and TMD was sufficient for envelope localization of GFP signal (Figure 6D). These results support the conclusion that the DEK5 TMD is necessary and sufficient for envelope localization.

Polyclonal α-DEK5 antibodies raised against the DEK5 TamB domain were used to determine native protein localization in maize (Figure 5C). The antibodies cross-react with a ~100 kDa polypeptide that co-purifies with wild-type chloroplasts as well as a larger band in total leaf protein (Supplemental Figure 9A). Pre-incubation of the α-DEK5 sera with the TamB recombinant antigen causes loss of the 100 kDa signal showing that this band is specific for the DEK5 protein despite migrating much faster than the full-length protein (Supplemental Figure 9B). Moreover, the specific α-DEK5 signal is reduced in *dek5* leaf extracts and the amount of DEK5 protein correlates with the strength of the mutant phenotype (Figure 6E). The severe *dek5-Briggs* and *dek5-N1339A* alleles have little to no detectable DEK5 protein as well as severely reduced outer envelope (Toc75, OEP80) and thylakoid (Cyt f) membrane proteins (Figure 6E). These data indicate that α-DEK5 reports DEK5 protein. Potentially, DEK5 is proteolytically processed in the cell or runs aberrantly in SDS-PAGE.

Sub-fractionation of the chloroplast to separate soluble, thylakoid membrane, inner envelope membrane (IEM), and outer envelope membrane (OEM) proteins showed that DEK5 co-fractionated with the envelope membranes and soluble proteins (Figure 6F). Chloroplast compartment specific protein markers show clean separation of soluble proteins from envelope membranes.

A dual-protease protection assay was used to determine the topology of DEK5 (Froehlich, 2011). Thermolysin only digests proteins on the outer face of the OEM. DEK5, 6PGDH, and OEP80 were all resistant to thermolysin when chloroplasts are intact (Figure 6G). Each of these proteins are sensitive to thermolysin when chloroplasts are lysed under hypotonic conditions.

By contrast, trypsin digests protein domains in the intermembrane space between the OEM and IEM. The OEM protein, OEP80, is sensitive to trypsin, while stromal 6PGDH is resistant (Figure 6G). DEK5 was partially digested by trypsin in five biological replicates (Figure 6G, Supplemental Figure 9C). All three proteins are fully digested in hypotonic chloroplast lysates. These results suggest that DEK5 protein is sensitive to trypsin and likely to face the intermembrane space between the OEM and IEM. This is a similar topology to TamB in *E. coli* but is contrasting to the stromal localization reported for rice SSG4 (Selkrig et al., 2012; Matsushima et al., 2014).

### Chloroplast envelope protein composition is altered in *dek5*

DEK5 envelope localization and intermembrane space topology supports the model that DEK5 has an orthologous function to TamB and may facilitate insertion of a subset of membrane proteins. Proteomics of purified chloroplast envelope fractions supports broad changes in envelope membrane protein composition (Figure 7A). This analysis identified 68 known chloroplast envelope protein homologs including 11 OEM proteins, 41 integral IEM proteins, 10 peripheral IEM proteins, and 6 envelope proteins with unknown locations. We compared envelope protein abundance by normalizing the MaxQuant intensity values for the 68 envelope proteins detected between *dek5* and normal samples (Supplemental Table 1). Nearly 1/3 of these proteins were reduced or absent in *dek5*. Among the 22 envelope proteins with reduced abundance, 91% are predicted to function in membrane transport.

**Figure 7.**
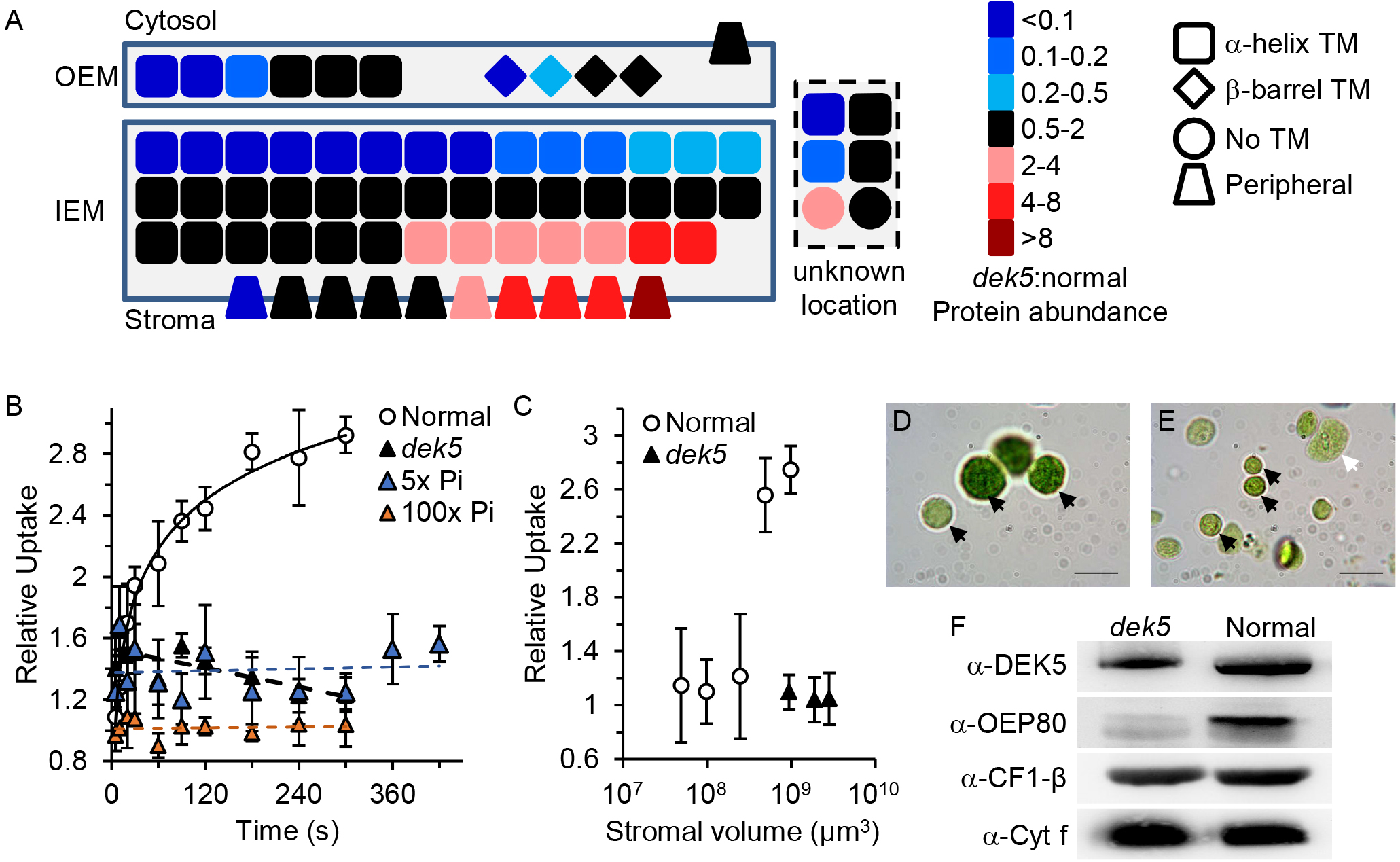
Defective *dek5* chloroplasts envelope membranes. (A) Heatmap of chloroplast envelope protein differences between *dek5* and normal siblings. Symbols indicate the type of transmembrane domain (TM) predicted for each protein. Proteins in the dotted box have not been localized specifically to the outer (OEM) or inner (IEM) envelope membranes. (B) Time course of ^32^Pi uptake in isolated chloroplasts. Mean and standard deviation of three biological replicates are plotted. (C) End-point ^32^Pi uptake of normal and *dek5-N961* chloroplast dilutions after 5 min incubation. Mean and standard deviation of three biological replicates are plotted. Total stromal volume was estimated based on the number of chloroplasts included in each assay. (D-E) Phase contrast micrographs of *dek5-N961* (D) and normal sibling (E) chloroplasts after a mock Pi uptake assay. Black arrows indicate intact chloroplasts. White arrows indicate broken chloroplasts. Scale bars are 10 μm. (F) Immunoblot analysis of marker proteins for purified *dek5-N961* and normal sibling chloroplasts. Protein was loaded on an equal OD_652_ basis.

Half of the OEM integral membrane proteins (5/10) were downregulated in *dek5* with the remaining OEM proteins being unchanged in relative abundance (Figure 7A). OEM proteins with reduced levels included the chloroplast protein targeting machinery, Toc34, Toc75 and Toc159, as well as OEP16 and OEP80. These results are consistent with immunoblots of *dek5* leaf protein showing that Toc75 and OEP80 are reduced (Figure 6E).

There were seven membrane proteins with increased relative abundance in the IEM. However, 14 IEM proteins have reduced relative levels in *dek5* including a Na^+^-dependent inorganic phosphate (Pi) transporter and two Pi/phosphoenolpyruvate translocators (Supplemental Table 1). Both α-helical and β-barrel transmembrane proteins are reduced in *dek5*, and we conclude that there are pleiotropic defects in envelope protein composition.

### Pi uptake is compromised in *dek5* chloroplasts

The reduced Pi transporter levels in *dek5* envelopes suggest that Pi transport may be affected. Pi uptake was assayed by incubating purified chloroplasts with ^32^Pi (Figure 7B). Normal chloroplasts reach equilibrium ^32^Pi levels between 3-5 min in 0.1 mM Pi. Mutant *dek5-N961* chloroplasts show no appreciable uptake over the 5 min incubation at 0.1 mM Pi. Higher Pi concentrations of 0.5 mM (5x) or 10 mM (100x) did not increase uptake to normal levels. A longer incubation time at 0.5 mM Pi also showed no appreciable uptake.

Pi accumulates in the stroma, and uptake is limited by total stromal volume in the assay (Fliege et al., 1978). A dilution series of normal sibling chloroplasts found that the detection limit of ^32^Pi uptake is >20% and <50% of the stromal volume included in the standard assay (Figure 7C). Although *dek5* stromal volume is much larger than in normal chloroplasts, *dek5* thylakoids account for approximately 10% more of the total chloroplast volume leading to a relative reduction in stroma. To normalize Pi uptake on stromal volume, we counted chloroplast number and estimated the average stroma volume using ImageJ (Supplementary Figure 5C). A dilution series from 1.7- to 5.8-fold larger stromal volume relative to the standard assay with normal chloroplasts showed no appreciable Pi uptake (Figure 7C). Based on the normal and *dek5* chloroplast dilution series, we expected to observe uptake if *dek5* membrane had as little as 17% of normal Pi uptake.

To control for chloroplast integrity throughout the assay, we completed Pi uptake assays without radioactive labeling and purified the chloroplasts in an isosmotic sorbitol solution. Phase contrast microscopy of the recovered chloroplasts showed that *dek5* chloroplasts were intact (Figure 7D-E). Immunoblots of purified chloroplasts loaded on an equal chlorophyll content showed equivalent levels of thylakoid membrane proteins, CF1-β and Cyt f, as expected from this normalization (Figure 7F). DEK5 and OEP80 were reduced relative to thylakoid membrane proteins consistent with reduced mutant chloroplast surface area and the relative reduction of OEP80 observed by proteomics. Together, these data argue that multiple membrane transporters and Pi transport are greatly reduced in *dek5* chloroplasts.

## DISCUSSION

Cloning of *dek5* revealed that the locus is orthologous to rice *ssg4* (Matsushima et al., 2014). However, we have come to different conclusions about the biochemical function of plant TamB proteins. Matsushima et al. (2014) suggested that SSG4 has a novel function in chloroplast division, while we propose that the DEK5/SSG4 proteins have analogous envelope biogenesis roles to bacterial TamB proteins.

A series of alleles with differing severity was critical in revising the model for plant TamB homologs. The rice *ssg4* locus is defined by a single mutant allele with a mis-sense mutation in the TamB domain (Matsushima et al., 2014). Compared to *dek5*, *ssg4* shows mild starch granule and chloroplast phenotypes with no ultrastructural changes observed in plastid membranes. By contrast, the Arabidopsis *embryo defective2410* locus also encodes a DEK5 ortholog, and the described T-DNA insertion alleles arrest seed development at the globular embryo stage (Tzafrir et al., 2004). The *dek5* allelic series shows that weak alleles express some DEK5 protein to give more normal development, while strong alleles lacking DEK5 protein cause severe seed defects and embryo lethality. Much of our phenotypic analysis focused on the mild *dek5-MS33* and *dek5-N961* alleles, which enable sufficient plastid development to observe a range of organelle defects. From these phenotype data, we conclude that *dek5* is required for envelope biogenesis and plastid metabolite transport.

### DEK5 is required for plastid metabolite transport

Multiple lines of evidence suggest that intracellular metabolite transport is disrupted in *dek5*. A primary function of endosperm amyloplasts is to accumulate starch. Endosperm starch levels are reduced in *dek5*. Sucrose is the major carbon source for endosperm starch synthesis, and *dek5* has increased levels of sucrose during seed development suggesting that carbon is not limiting for starch production. For starch synthesis, sucrose is cleaved and converted to ADP-glucose in the cytoplasm of endosperm cells (Hannah, 1997). The BT1 inner envelope membrane transporter selectively transports ADP-glucose into the amyloplast (Shannon et al., 1998). Like *bt1*, *dek5* mutants have higher levels of ADP-glucose suggesting reduced transport of ADP-glucose into amyloplasts may be the primary cause for increased sugar levels in *dek5* (Tobias et al, 1992; Shannon et al., 1996). This inference is further supported by the nearly identical starch branch chain lengths in *dek5* and normal endosperm starch, which indicates starch polymerization and debranching is normal in *dek5*. Moreover, *dek5* starch granule size is increased showing robust starch synthesis when sufficient substrate is available.

There is also evidence for solute transport defects in *dek5* leaves. Maize is a C4 plant that fixes CO_2_ in mesophyll chloroplasts and shuttles malate into bundle sheath chloroplasts for starch synthesis (Majeran et al., 2005; Weise et al., 2011). TEM of *dek5* bundle sheath chloroplasts found few starch granules but extensive thylakoid vesicular structures consistent with an osmotic imbalance between the chloroplast and cytosol. Moreover, *dek5* mesophyll chloroplasts accumulate starch, which would be consistent with reduced flux of malate from the mesophyll to bundle sheath.

Proteomics analysis revealed that 33% of envelope proteins are reduced >2-fold in *dek5*. The majority of the affected proteins function in metabolite and ion transport. Moreover, we were not able to detect Pi uptake in *dek5* chloroplasts *in vitro*. The sensitivity of the assay indicates that *dek5* chloroplasts have at least an 80% reduction in Pi transport. These data support a role for DEK5 in envelope protein accumulation and metabolite transport.

### DEK5 is required for chloroplast membrane biogenesis

The proteomics and Pi uptake data are consistent with *dek5* chloroplast shape and membrane ultrastructure. Mutant chloroplasts have a more spherical shape than normal. Consequently, envelope surface area is reduced in *dek5* relative to chloroplast and thylakoid volume. Arabidopsis chloroplast division mutants such as *arc3*, *arc5, arc6*, and *acr8*, have fewer and larger chloroplasts (Pyke and Leech, 1992; Pyke et al., 1994; Robertson et al., 1996; Shimada et al., 2004; Maple et al., 2007). However, the enlarged *arc* chloroplasts elongate instead of becoming spherical, which maintains envelope surface area to volume ratios closer to wild-type. The *arc* mutants show that defects in the plastid division machinery does not limit envelope membrane biogenesis.

The *dek5* chloroplasts also expand the peripheral reticulum, which is a tubular network derived from the inner envelope (Wise, 2007). Potentially, this ultrastructural change indicates differential development of inner and outer envelope membranes. There is some similarity between the *dek5* envelope phenotypes and a mutation in a TamB homolog, morC, in *Aggregatibacter actinomycetemcomitans.* The morC mutation reduces outer membrane relative to inner membrane resulting in a smooth outer surface when wild-type cells have convoluted, rugose outer membranes (Gallant et al., 2008; Azari et al., 2013). Like *dek5* chloroplasts, morC mutants have a more spherical cell morphology suggesting that loss of TamB function can lead to similar membrane imbalance in both bacteria and chloroplasts.

Envelope proteomics showed that *dek5* has a different relative protein composition in both envelope membranes. Similar proteomics studies have not been completed in bacterial TamB mutants, but mutants in *E. coli*, *A. actinomycetemcomitans*, and *Deinococcus radiodurans* all disrupt localization of a subset of outer membrane proteins (Selkrig et al., 2012; Azari et al., 2013; Shen et al., 2014; Yu et al., 2017). In the *dek5* outer envelope, half of the membrane proteins have reduced relative levels, while the inner envelope has a subset of proteins with increased levels. This further argues that the outer envelope is more compromised relative to the inner envelope.

Thylakoids can also be disrupted in *dek5*, but the loss of thylakoid organization is likely due to defects in envelope protein composition. Both proteomics and immunoblotting found that subunits of the chloroplast protein import machinery are reduced in *dek5* envelopes. Toc34, Toc75, and Toc159 are the core translocation complex (Schleiff et al., 2003). All three of these proteins have reduced levels in *dek5* envelopes. Based on immunoblots, Toc75 levels correlate with the amount of DEK5 protein in the *dek5* allelic series. Toc75 is a β-barrel integral membrane protein that is the channel for translocation of nuclear-encoded proteins into the chloroplast (reviewed in Bölter and Soll, 2016). Loss of Toc75 in Arabidopsis causes early embryo arrest due to a loss of import of nuclear-encoded proteins into the chloroplast (Schleiff et al., 2003; Hust and Gutensohn, 2006). The reduced import machinery in *dek5* is expected to limit chloroplast protein accumulation to cause the defects observed in thylakoid membranes.

Reduced levels of Toc75 may be a more primary consequence of the *dek5* mutation. The function of TamB in *E. coli* is to promote integration of β-barrel membrane autotransporters (Selkrig et al., 2012; Shen et al., 2014). In *E. coli*, TamB interacts with TamA as well as the β-barrel assembly module (BAM) subunits (Babu et al., 2018). TamB is found in more prokaryotes than TamA, but TamB has a conserved function in outer membrane biogenesis as demonstrated in *D. radiodurans* (Yu et al., 2017). Thus, it is likely that DEK5 plays a direct role in targeting β-barrel membrane proteins to the outer envelope.

### DEK5 topology is analogous to TamB

Biochemical analysis of DEK5 topology further supports the model that plant TamB proteins have an analogous function with bacterial homologs. TamB is an inner plasma membrane protein with an N-terminal transmembrane domain and the C-terminal TamB domain that bridges the periplasmic space to interact with TamA or the BAM complex (Selkrig et al., 2012; Shen et al., 2014). Plant TamB proteins are much bigger than homologous proteins in proteobacteria. Matsushima et al (2014) proposed that SSG4 has a distinct biochemical function from TamB, in part, based on these protein sequence differences. However, Gram-negative bacteria have a 2-6 nm periplasm, while cyanobacteria and chloroplasts have thicker periplasm and intermembrane spaces of 10-35 nm (Hoiczyk and Hansel, 2000). In *E. coli*, TamB is integrated into the plasma membrane, bridges the periplasm, and physically associates with TamA in the outer membrane (Shen et al., 2014). Plant DEK5 proteins would need to be larger to bridge the intermembrane space of the plastid.

When fused to GFP, the N-terminal region of DEK5 encompasses chlorophyll fluorescence in a pattern described for known envelope proteins (Okawa et al., 2008; Singh et al., 2008; Kasmati et al., 2011). By contrast, the rice SSG4N-GFP fusion localized to the stroma in both *N. benthamiana* and rice (Matsushima et al., 2014). Potentially, differences in truncation of DEK5 and SSG4 proteins used for the fusions led to different suborganellar localization. The SSG4N-GFP encodes the N-terminal 213 amino acids of the SSG4 protein, which disrupts a conserved protein sequence found in all plant DEK5 homologs (Supplemental Figure 8). Protein sequence targeting determinants have only been identified for a few single-pass transmembrane inner envelope proteins (Viana et al., 2010; Froehlich and Keegstra, 2011). It is possible that the partial truncation of the homology domain in the SSG4N-GFP fusion causes defective targeting to the stroma.

Importantly, Matsushima et al (2014) used a single GFP fusion protein as the sole evidence of stromal localization of SSG4. We also raised antibodies that cross-react with the C-terminal region of DEK5. The native DEK5 protein co-purifies with envelope membranes, is partially sensitive to trypsin digestion, and is found in the soluble chloroplast protein fraction after hypotonic lysis. Hypotonic lysis of chloroplasts is expected to mix soluble envelope intermembrane space proteins with stroma proteins. Few soluble intermembrane space proteins have been identified, but these do co-purify with the soluble fraction (Hirohashi et al., 2001; Bionda et al., 2016). Thus, the presence of the native DEK5 protein in the envelope and soluble fractions as well as trypsin sensitivity argues that DEK5 is an intermembrane space protein associated with both outer and inner envelopes. This is the same topology as the *E. coli* TamB protein (Selkrig et al., 2012; Shen et al., 2014).

### A revised model for plant TamB homologs

Combining the experimentally determined topology of the DEK5 protein with the reduced metabolite transport, reduced envelope protein levels, and reduced outer envelope membrane in *dek5* chloroplasts leads to the conclusion that DEK5 functions in chloroplast envelope biogenesis. Based on TamB function, we propose that DEK5, SSG4, and EMB2410 have conserved roles in targeting a subset of β-barrel proteins to the outer envelope. The proteomics experiment identified Toc75 and OEP80 as likely targets. OEP37 levels were just above the 2-fold reduction cut-off in the proteomics analysis and may also be a DEK5 substrate. The correlation between the levels of chloroplast import machinery and the severity of *dek5* mutant alleles argues that severe alleles display extreme pleiotropy with plastids being unable to support import of nuclear-encoded proteins. Weak alleles that support protein import will also retain sufficient DEK5 activity to support β-barrel protein insertion. Consequently, direct proof of the proposed function in outer envelope protein insertion will require biochemical methods to block DEK5 action.

## METHODS

### Plant materials

All maize plants were grown during April-July or August-December at the University of Florida Institute of Food and Agricultural Sciences Plant Science Research and Education Unit in Citra, Florida. The *dek5* and normal siblings were germinated in a greenhouse with Metro-Mix 300 (Scotts-Sierra. Marysville, OH). The *dek5-N874A*, *dek5-N961,* and *dek5-N1339A* were originally isolated by Neuffer and Sheridan (1980) from EMS mutagenesis. Scanlon et al. (1994) isolated *dek5-PS25* and *dek5-MS33* from *Mutator* transposon-tagging populations. The *dek5-Briggs* allele is a spontaneous allele (www.maizegdb.org). Stocks for *dek5* alleles were ordered from the Maize Genetics COOP Stock Center. Each *dek5* allele was crossed with B73, W22, or Mo17 inbred lines five times to develop BC_4_ introgression stocks to characterize phenotypes. The *dek5-umu1* (mu1021358) and *dek5-umu3* (mu1047944) UniformMu alleles were maintained in the W22 inbred background. We confirmed all stocks were *dek5* alleles by genetic complementation tests using reciprocal crosses between each stock and *dek5-PS25*. First ears were used for reciprocal crosses. Second ears of both male parents and female parents in the crosses were self-pollinated to determine plant genotypes.

### Endosperm composition analysis

Endosperm composition was determined from dissected tissues. Developing endosperm was flash frozen in liquid nitrogen and stored at -80°C. Frozen endosperm tissues were cracked into small pieces with mortar and pestle, lyophilized for 3 days, and ground to a fine powder with a bead mill. For mature dry kernels, endosperm was isolated by cutting the kernel in the sagittal plane and cutting away the embryo with a utility knife. The pericarp was removed by scraping the surface of the mature dry endosperm with a razor blade. The remaining endosperm tissue was ground to a fine powder with a bead mill.

Zein and non-zein protein were extracted from the dissected, dry endosperm tissue as described by Wu and Messing (2012). Protein extracts were loaded on equal dry weight basis for SDS-PAGE and stained with Commassie Blue R-250. Endosperm starch was measured from developing kernels using the Megazyme Total Starch Assay Kit (AA/AMG). Sucrose, glucose, and maltose were measured with the Megazyme Maltose/Sucrose/D-Glucose Assay Kit (K-MASUG). ADP-Glucose was measured with a glycerol-3-phosphate oxidase/glycerol-3-phosphate dehydrogenase cycling assay as described (Gibon et al., 2002).

### Starch analysis

Developing kernels were collected and fixed in FAA solution (3.7% formaldehyde, 5% glacial acetic acid, and 45% ethanol) at 4°C overnight. Kernels were dehydrated in an ethanol series and kept in 100% ethanol. Additional sample preparation and SEM was performed in Interdisciplinary Center for Biotechnology Research Electron Microscopy core at the University of Florida, Gainesville, FL. For particle size analysis, mature endosperm tissue was dissected and ground into fine powder with a bead mill. Endosperm powder (0.1 g) was mixed with 1 mL of ethanol and vortexed immediately prior to measurement with a Beckman/Coulter LS 13 320 laser diffraction particle size analyzer. Linear chain length distributions of endosperm starch was determined using fluorophore-assisted carbohydrate electrophoresis (FACE) with a Beckman P/ACE capillary electrophoresis system as described (O’Shea et al., 1998; Dinges et al., 2003).

### Protoplast analysis

Maize protoplasts were prepared essentially as described with some modifications (Yoo et al., 2007). Kernels were planted in a greenhouse. After 10 d, 0.5 g seedling leaves were harvested, chopped with a razor blade, and digested in enzyme solution at 28°C for 4-5 h with a gentle shaking at 60 rpm in the dark. The enzyme solution consisted of 1.5% (w/v) cellulase R10, 0.5% (w/v) macerozyme R10 in 0.4 M mannitol, 20 mM KCl, 10 mM CaCl_2_, 5 mM β-mercaptoethanol, 0.1% BSA and 20 mM MES (pH 5.7). Cell debris was filtered using 35 μm nylon mesh, and collected protoplasts were suspended in W5 solution (2 mM MES pH 5.7, 154 mM NaCl, 125 mM CaCl_2_, 5 mM KCl) and kept on ice for observation. Protoplast isolation at high osmotic concentration replaced 0.4 M mannitol with 0.8 M mannitol in all solutions. Protoplast yield was determined by counting intact protoplasts in a hemocytometer with light microscope. Light microscopy imaging used an AMScope digital camera.

Chloroplast number within each cell was determined with Z stack images of individual protoplast cells using a Leica TCS SP5 laser scanning confocal laser scanning microscope (Leica Microsystems). Chlorophyll was excited at 488 nm and detected with an emission band of 650-700 nm. A 3D image was reconstructed for each cell and manually scored for the number of chloroplasts. Chloroplast cross section surface areas and volume were measured using image J (Schindelin et al., 2012).

### Leaf pigment extraction and measurement

Fresh seedling leaf tissue (0.5 g) from both *dek5* mutant and normal siblings was harvested and chopped with a razor blade. The leaf tissue was extracted in 15 mL of 96% ethanol with gentle agitation for 12 h at room temperature in aluminum foil wrapped bottles. Ethanol extracts were separated from tissue debris with filter paper. The tissue was rinsed with 96% ethanol to completely extract residual pigment. The extract and rinse were combined and brought to 25 mL final volume in 96% ethanol. Chlorophyll-a, chlorophyll-b, and carotenoid absorbance were measured at 665, 649, and 470 nm using a spectrophotometer. Pigment content was calculated according to Hartmut (1983).

### Immunoblotting

Chloroplast protein or total seedling proteins were separated by 10% SDS-PAGE, and transferred to nitrocellulose membrane using a Bio-Rad Trans-Blot SD semi-dry transfer cell at 15 V for 30 min. Membrane was first blocked with blocking solution containing 3% BSA and 0.2% TWEEN 20 with PBS (pH 7.4) at room temperature for 30 min, then incubated with primary antibody at 1:2,000 dilution in blocking solution for 2 h at room temperature or overnight at 4°C. Membranes were washed four times for 5 min using PBS with 0.2% TWEEN 20. Primary antibody signal was detected after incubating goat anti-rabbit secondary antibody (Sigma) at 1:1,000 dilution in blocking solution for 2 h at room temperature followed by four 5 min washes using PBS with 0.2% TWEEN 20. Chemiluminescent was produced by applying 1 mL ECL substrate (Thermo Scientific Pierce) per gel membrane and imaging with a FOTODYNE FOTO Luminary/FX digital imaging system.

### Leaf transmission electron microscopy (TEM)

Seedling leaf tissues were harvested 10 DAS in a growth chamber. Sample fixation, sectioning and TEM were completed by Electron Microscopy Services, LLC., as described (Turgeon and Medville, 2004). Briefly, leaf samples were fixed for 4 h at 4°C in 2% (w/v) paraformaldehyde and 2.5% (v/v) glutaraldehyde in 70 mM sodium cacodylate buffer, pH 6.8, followed by overnight incubation in 1% (w/v) osmium tetroxide in fixation buffer. Fixed tissues were subsequently dehydrated and embedded in resin. Ultrathin sections were stained with uranyl acetate and lead citrate for TEM observations.

Chloroplast area, long axis, and short axis were measured using ImageJ (Schindelin et al., 2012). Stroma and thylakoid area were estimated by first determining the average auto threshold value for all TEM images. The average threshold value was applied to all images to measure thylakoid area, which was subtracted from chloroplast area to estimate stroma area. Volume and surface area calculations were calculated assuming the short axis represents the height and width of an ellipsoid, while the long axis represents the length. Thylakoid and stroma volumes were calculated for each chloroplast based on the relative fraction of the total chloroplast area.

### Molecular identification of the *dek5* locus

An F_2_ mapping population was generated by self-pollinating F_1_ plants from a cross of the Mo17 inbred and *dek5-PS25/+*. Genomic DNA was extracted from individual F_2_ mutant kernels and analyzed with simple sequence repeat (SSR) and insertion-deletion (InDel) PCR markers as described (Settles et al., 2014). Supplemental table 4 gives primer sequences of the markers used for linkage analysis. For the transposon flanking sequence experiment, homozygous *dek5-PS25* and *dek5-MS33* mutant genomic DNA was isolated from pools of six F_2_ mutant kernels. *Mu* flanking sequences were amplified, purified, and adapted to the Illumina sequencing platform as described (McCarty et al., 2013). Reads were quality filtered and mapped to the B73_v3 genome assembly using the UniformMu informatics pipeline (McCarty et al., 2013).

### RNA Isolation and RT-PCR

Total RNA was isolated using Trizol (Invitrogen) from the following B73 maize tissues: leaves, root, silks, tassel, and developing kernels at different stages. Approximately 1 mg of total RNA was reverse transcribed with either ThermoScript Reverse Transcriptase (Invitrogen) for Dek5 cDNA cloning or SuperScript III First-Strand Synthesis System (Invitrogen) for semi-quantitative RT-PCR. Both oligo (dT) and random hexamer primers were used for first-strand synthesis to ensure full-length coverage of the *Dek5* mRNA. The *Dek5* cDNA was amplified with Phusion high-fidelity DNA polymerase (New England Biolabs) and cloned into pENTR /D-TOPO Vector (Invitrogen) as three overlapping fragments: *Dek5*-FN (1802 bp), *Dek5*-FM (2330 bp), and *Dek5*-FC (3006 bp). The full length *Dek5* cDNA was assembled from the sequence of these clones. Specific PCR primers are listed in Supplemental table 4.

### Phylogenetic Analysis

Protein sequences were aligned using ClustalW (Larkin et al., 2007). The phylogenetic tree was produced with MEGA7 using the neighbor-joining method with 2,000 bootstrap replications and the Poisson substitution model (Hedges, 1992). The DEK5 homolog sequences used for phylogenetic analysis were: *Zea mays* (MF034077); *Sorghum bicolor* (XP_002457235.1), *Glycine max* (XP_003527803.1), *Oryza sativa Japonica* (XP_015619226.1), *Setaria italic* (XP_004971719.1), *Fragaria vesca subsp. Vesca* (XP_011459147.1), *Vitis vinifera* (XP_010648561.1), *Arabidopsis thaliana* (NP_180137.3), *Theobroma cacao* (EOY31352.1), *Amborella trichopoda* (XP_006844393.1), *Selaginella moellendorffii* (XP_002982313.1), *Ostreococcus tauri* (XP_003079877.1), *Nostoc punctiforme* (WP_012412249.1), *Cyanobacterium PCC 7702* (WP_017320409.1), *Pseudomonas stutzeri* (WP_003289734.1), *Methylobacterium sp. Leaf91* (WP_082490488.1), *Rhizobium leguminosarum* (WP_003543422.1), *Arhodomonas aquaeolei* (WP_018716326.1), *Escherichia coli* (WP_040089969.1).

### Subcellular localization of DEK5

The DEK5N-GFP fusion protein was constructed by amplifying the initial 1,374 bp of the *Dek5* ORF from B73 seedling cDNA using Phusion high-fidelity DNA polymerase (New England Biolabs) with the *Dek5*N-L and *Dek5*N-R primers (Supplemental table 4). The N-terminal ORF product was cloned into pENTR vector (Invitrogen) and recombined into the binary vector, pB7FWG2, to generate a C-terminal GFP fusion (Karimi et al., 2002; Karimi et al., 2007). The TMD deletion was generated with the Q5 Site-Directed Mutagenesis Kit (New England Biolabs) to delete the peptide sequence, LFLVRCSVFAAVVSVAGALSW, in-frame from the DEK5N clone and then recombined into pB7FWG2. The binary constructs were transformed into *Agrobacterium tumefaciens* strain GV3101 by the freeze-thaw method (Weigel and Glazebrook, 2006). Each construct was transferred to *N. benthamiana* leaves via agroinfiltration as described (van Esse, 2012) with some modifications. Before infiltration, the agrobacteria were suspended in buffer containing 10 mM MgCl_2_, 10 mM MES and 200 μM acetosyringone and adjusted to OD_600_ of 0.4. The *N. benthamiana* plants were grown in a growth chamber at 23°C with 16/8 h day/night light cycle. Leaves from 4-5 weeks-old plants were infiltrated and examined by epifluorescence microscopy between 40-50 h after infiltration. Representative images were captured using a Leica TCS SP5 laser scanning confocal laser scanning microscope (Leica Microsystems). GFP was excited at 488 nm and detected with an emission band of 500-530 nm. Chlorophyll was detected with an emission band of 650-700 nm.

### DEK5 antibodies

The coding sequence for the DEK5 TamB domain was amplified with Phusion high fidelity DNA polymerase from B73 cDNA with the DUF490-L and DUF490-R primers (Supplemental table 4). The PCR product was digested with *Nco*I and *Not*I, purified, and cloned into the pMAL-c5X vector (New England Biolabs). Recombinant protein was expressed in the *E. coli* Top10 strain by inducing 12 L of culture with 10 mM IPTG for 1 h at 30°C. Cells were sonicated using a 5 s/5 s pulse/pause cycle for a total of 3 min in an ice-water bath. Recombinant protein was purified using amylose resin according to the manufacturer’s instructions (New England Biolabs). The recombinant protein was concentrated in Amicon Ultra-15 Centrifugal Filters (EMD Millipore) and 2.5 mg of purified recombinant protein was sent to ThermoScientific Pierce for rabbit polyclonal antibody production.

### Chloroplast isolation

Normal chloroplasts were isolated from normal siblings essentially as described (Voelker and Barkan, 1995). Leaf tissue from 7 day-old plants was cut and homogenized in grinding buffer (50 mM HEPES/KOH, pH 7.5, 0.33 M sorbitol, 1 mM MgCl_2_, 1 mM MnCl_2_, 2 mM EDTA, 5 mM Na-ascorbate, and 1% BSA) using a Warring blender with five bursts of ~5 s at max speed. The homogenate was filtered through 2 layers of Miracloth, and the filtrate was centrifuged at 3,000 g for 5 min in a swinging bucket rotor at 4°C. The organelle pellet was gently suspended in 5 mL grinding buffer and layered onto a 35%/75% (8 mL/16 mL) Percoll step gradient. The gradient was centrifuged at 6,000 g for 15 min and the lower, intact chloroplast, band was recovered. Intact chloroplasts were washed with import buffer (0.33 M sorbitol, 50 mM HEPES-KOH, pH 8.0) and pelleted by centrifugation at 1,000 g for 5 min. Pelleted chloroplasts were suspended in import buffer and adjusted to 0.5-1.0 mg chlorophyll per mL based on OD_652_.

Mutant *dek5* chloroplasts were purified from protoplasts. Isolated *dek5* mutant protoplasts were washed and suspended in a protoplast hypotonic lysis buffer of 0.7 M sorbitol supplemented with 50 mM HEPES-KOH (pH 7.5). The cell lysate was passed through a 15 μm nylon mesh filter (Tisch Scientific) to further lyse remaining intact protoplasts. Released *dek5* mutant chloroplasts were purified in a 25%/75% Percoll step gradient centrifuged at 6,000 g for 15 min, and the lower, intact chloroplast band, was recovered. Isolated *dek5* chloroplasts were washed with 0.7 M sorbitol and 50 mM HEPES-KOH (pH 7.5), pelleted and adjusted to 0.5-1 mg/mL chlorophyll.

### Chloroplast fractionation

Chloroplasts were fractionated as described by Smith et al. (2002) with some modifications. Briefly, intact isolated chloroplasts were pelleted from import buffer and resuspended to 1-2 mg/mL chlorophyll in TE buffer (10 mM Tricine, 2 mM EDTA, pH 7.5) with 0.6 M sucrose. Chloroplasts were lysed with two cycles of snap freezing and thawing and then diluted with TE buffer containing 0.2 M sucrose and protease inhibitor cocktail (Sigma, P9599). Lysed chloroplasts were centrifuged at 45,000 g for 60 min to collect the crude envelope membranes. The soluble supernatant was collected and stored at -80°C for further analysis. The membrane pellet was resuspended in TE buffer containing 0.2 M sucrose to a concentration of 2 to 3 mg/mL chlorophyll, and 1 mL of the membrane suspension was layered onto the top of a sucrose gradient. The gradient consisted of 1.5 mL of 1 M sucrose, 1 mL of 0.8 M sucrose, and 1 mL of 0.46 M sucrose in a 5 mL polyallomer tube. The gradient was centrifuged for 1.5 h at 270,000 g at 4°C using low acceleration and deceleration rates. The yellow-green band from the 0.8 to 1 M sucrose interface was collected as the inner envelope membrane fraction. The light yellow band from the 0.46 to 0.8 M sucrose interface was collected as the outer envelope membrane faction. The green thylakoid pellet was resuspended in TE. All membrane fractions were washed using 3 to 5 volumes of TE buffer supplemented with protease inhibitor and centrifuged for 1.5 h at 270,000 g at 4°C.

### Protease protection assay

The dual protease protection assay was done according to Froehlich (2011) with some modifications. Intact chloroplasts were isolated and suspended in import buffer (0.33 M sorbitol, 50 mM HEPES/KOH, pH 8.0) to a concentration of 1 mg/mL chlorophyll. Intact chloroplasts (150 μL) were incubated with either 25μL thermolysin or trypsin (1 mg/mL) for 50 min on ice. Thermolysin was stopped using 10 mM EDTA and kept on ice for 5 min. The trypsin digestion was stopped using 50 μL of protease inhibitors containing 1 mg/mL trypsin inhibitor (Sigma), 10 μM/mL leupeptin and 10 mM phenylmethylsulfonyl fluoride (PMSF). Intact chloroplasts were recovered using 30% Percoll cushion in import buffer and centrifuged at 1,500 g for 5 min. Pelleted intact chloroplasts were washed and resuspended in import buffer for further analysis.

### Proteome analysis of the *dek5* chloroplast envelope and its normal siblings

To collect enough chloroplast envelope proteins for proteome analysis, six independent chloroplast preps from *dek5* and four independent chloroplast preps from normal siblings were pooled separately and resuspended to 1-2 mg/mL chlorophyll in suspension buffer (0.6 M sucrose, 10 mM Tricine, 2 mM EDTA, pH 7.5). Both the *dek5* and normal chloroplasts were lysed with two cycles of snap freezing and thawing and then diluted with TE buffer containing 0.2 M sucrose and protease inhibitor cocktail. A Dounce homogenizer was used to further rupture the *dek5* chloroplasts. Lysed chloroplasts were centrifuged at 4,000 g for 15 min to remove most of the thylakoid membranes. The supernatant was transferred to a new tube and centrifuged at 40,000 g for 60 min to collect crude envelope membrane fraction. Enriched chloroplast envelope proteins were solubilized in 8M urea, 100 mM Tris-HCL pH 7, and 5 mM Tris-(2-Carboxyethyl) phosphine, Hydrochloride (TCEP). Proteins were then subjected to four rounds of acetone precipitation with probe sonication between each round. Pelleted protein was solubilized in UA buffer (8M urea, 50mM Tris-HCL pH 7, and 5 mM TCEP) and processed using the FASP method (Wisniewski et al., 2009; Song et al., 2017) with microcon YM-30 centrifugal filters (Millipore). Proteins were digested on the filter overnight with 1 μg trypsin (Roche) at 37°C. Proteins were then further digested with 0.1 μg of trypsin and 0.1 μg of LysC (Wako) for 2 h. Recovered peptides were acidified to a pH of ~2-3 with formic acid and desalted with 50 mg Sep-Pak C18 cartridges (Waters). Eluted peptides were dried in a vacuum centrifuge, resuspended in 0.1% formic acid, and quantified using the Pierce BCA Protein assay kit.

One μg of peptides were separated on a 20 cm nanospray column, which was pulled and packed in-house with 2.5 μm C18 (Waters), using an acetonitrile gradient of 5-30% for 120 min, 30-80% for 25 min, and 0% for 5 min (150 min total) that was delivered via an Agilent 1260 quaternary HPLC at a flow rate of ~500 nL/min. The HPLC system was coupled with a Thermo Scientific Q-Exactive Plus high-resolution quadrupole Orbitrap mass spectrometer using a custom fabricated nano-spray source. Data dependent acquisition was obtained using Xcalibur 4.0 software in positive ion mode with a spray voltage of 2.00 kV, and RF of 60, and a capillary temperature of 275°C. MS1 spectra were measured at a resolution of 70,000 with an automatic gain control (AGC) of 3 x 10^6^, a maximum ion time of 100 ms, and a mass range of 400-2000 m/z. Up to 15 MS2, with a charge state of 2 to 4, were triggered at a resolution of 17,500, an AGC of 1 x 10^5^ with a maximum ion time of 50 ms, a 1.5 m/z isolation window, and a normalized collision energy of 28. MS1 that triggered MS2 scans were dynamically excluded for 25 seconds. We performed two runs for each sample to generate two technical replicates.

The raw data were searched against the B73 RefGen_v2 5b Filtered Gene Set using MaxQuant version 1.5.8.3 (Tyanova et al., 2016). Methionine oxidation and protein N-terminal acetylation were set as variable modifications. The digestion parameters were set to “specific” for “Trypsin/P;LysC” with a maximum of 2 missed cleavages. The match between runs feature was turned off. Default settings were used for the remaining parameters including a peptide spectrum match and protein false discovery rate (FDR) of 0.01, which was determined using a reverse decoy database.

The *dek5* mutant and normal sibling envelope protein extracts had different levels of non-envelope contaminants. To compare relative levels of envelope proteins, we curated a set of 96 envelope proteins that were validated experimentally. The MaxQuant intensity for the 85 envelope peptides within this subset found in at least one technical replicate were normalized relative to one normal technical replicate. Proteins identified in a single technical replicate with a normalized MaxQuant intensity value <1×10^7^ were removed to leave 68 proteins. Relative fold change between *dek5* and normal sibling samples were calculated from the average of the two technical replicates or the single MaxQuant intensity value if the protein was detected only once. Transmembrane domains for envelope proteins were predicted using TMHMM Server V2.0 (http://www.cbs.dtu.dk/services/TMHMM/), and envelope localization was annotated using the Plant Proteome Database (http://ppdb.tc.cornell.edu).

### ^32^Pi chloroplast uptake assay

Uptake of ^32^Pi into isolated chloroplasts was assayed as described with some modifications (Fliege et al., 1978). Purified, intact chloroplasts from B73 and *dek5-N961* mutants were adjusted in concentration to OD_652_ of 0.2 and 90 μL (approximately 2.3× 10^7^ normal and 4.6× 10^6^ *dek5*) chloroplasts were mixed with 10 μL 1 mM KH_2_PO_4_ (pH 7.5). KH_2_PO_4_ was prepared in a 19:1 ratio of unlabeled to radiolabeled 37 MBq/mM KH_2_^32^PO_4_ (PerkinElmer). Chloroplasts were incubated in a time course from 5 to 300 s. Phosphate uptake was stopped by adding 2 mM 4,4′-Diisothiocyanatostilbene-2,2′-disulfonic acid (DIDS, Sigma) to the reaction. Relative uptake was calculated from background radiolabel as determined by pre-incubating chloroplasts with DIDS for 5 min before incubating with ^32^Pi for 300 s. Intact normal chloroplasts with radiolabel were re-purified by transferring 100 μL of the chloroplast suspension to a centrifuge tube containing a lower layer of 30 μL of 10% perchloric acid and an upper layer of 70 μL AP 150 silicone oil (Wacker). The density of *dek5* mutant chloroplasts was lower than normal and intact chloroplasts were purified using a 4:1 mixture of AP 150 and AR 20 silicone oil (Sigma). The radiolabeled chloroplast suspension was overlaid onto the silicone oil and then centrifuged for 30 s at 14,000 g. The radiolabel associated with intact chloroplasts was assayed by sampling 15 μL of the 10% perchloric acid layer for scintillation counting using a Beckman LS6500SE scintillation counter.

Detection limits were determined after a 300 s incubation with normal chloroplasts diluted from 5-50% of an OD_652_ of 0.2. Mutant chloroplasts were concentrated by centrifugation of 2- and 4-fold larger volumes at 500 g for 1 min followed by resuspension in 90 μL of 0.7 M sorbitol, 50 mM HEPES-KOH (pH 7.5). Stromal volume was estimated by counting intact chloroplasts using a hemocytometer with a light microscope and multiplying by the average stromal volume from TEM.

Assay effects on chloroplast integrity was determined by incubating chloroplasts with 0.1 mM KH_2_PO_4_ (pH 7.5) for 300 s. Chloroplasts were re-purified through silicone oil with a bottom layer of 30 μL of 50 mM HEPES-KOH (pH 7.5) and 0.3 M or 0.7 M sorbitol for normal and *dek5* mutants, respectively. Pelleted, intact chloroplasts were washed twice with the sorbitol buffer to remove residual silicone oil, observed under a phase contrast microscope, and imaged with an AmScope digital camera.

## Acknowledgements

We thank Richard Medville at the Electron Microscopy Services (Colorado Springs, CO 80920) for performing the TEM analysis. We thank Dr. Antje von Schaewen at the University of Münster for providing the 6PGDH antibody, Dr. Carole Dabney-Smith at Miami University for providing the Toc75 antibody, Dr. Kentaro Inoue and Philip Day at UC Davis for providing the OEP80 antibody. We also thank Dr. Curtis Hannah and Dr. Ken Cline at the University of Florida for their intellectual input. This work was supported by the National Institute of Food and Agriculture (award 2010-04228), the China Scholarship Council, and the Vasil-Monsanto Endowment.

## Author contributions

J.Z. and A.M.S designed the research, analyzed data, and wrote the manuscript; J.Z. performed most of the experiments; S.W. performed the Mu-Seq experiment and D.R.M. analyzed the data; S.K.B. performed the ADP-glucose measurement; G.S. and J.W. performed the chloroplast envelope proteomic analysis. A.M. performed the amylopectin chain distribution experiment.

## Supplemental Data

**Supplemental Figure 1.** Kernel and seedling phenotypes of *dek5*.

**Supplemental Figure 2.** Composition phenotypes of *dek5* kernels.

**Supplemental Figure 3.** Genetic background effects on *dek5*.

**Supplemental Figure 4.** Range of chloroplast defects in *dek5*.

**Supplemental Figure 5.** Chloroplast traits based on TEM measurements.

**Supplemental Figure 6.** Molecular cloning of *Dek5*.

**Supplemental Figure 7.** Phylogenic analysis of DEK5 orthologous proteins.

**Supplemental Figure 8.** Multiple sequence alignment of N-terminal DEK5 protein sequences.

**Supplemental Figure 9.** DEK5 antibody control experiments.

**Supplemental Table 1.** Kernel composition analysis

**Supplemental Table 2.** Seedling leaf photosynthetic pigments

**Supplemental Table 3.** Proteomic analysis of chloroplast envelope proteins.

**Supplemental Table 4.** Primers used in this study.

**Supplemental Movie 1.** Z-stack images of chlorophyll autoflorescence in normal and *dek5* protoplasts showing lack of chlorophyll in the interior of some *dek5* chloroplasts.

